# Nature Notes: Spatiotemporal variation in the competitive environment, with implications for how climate change may affect a species with parental care

**DOI:** 10.1101/2022.12.20.521241

**Authors:** Ahva L. Potticary, Hans W. Otto, Joseph V. McHugh, Allen J. Moore

## Abstract

Burying beetles of the genus *Nicrophorus* have become a model for studying the evolution of complex parental care in a laboratory. *Nicrophorus* species depend on small vertebrate carcasses to breed, which they process and provision to their begging offspring. However, vertebrate carcasses are highly sought after by a wide variety of species and so competition is expected to be critical to the evolution of parental care. Despite this, the competitive environment for *Nicrophorus* is rarely characterized in the wild and remains a missing factor in laboratory studies. Here, we performed a systematic sampling of *Nicrophorus orbicollis* living near the southern extent of their range at Whitehall Forest in Clarke County, Georgia, USA. We determined the density of *N. orbicollis* and other necrophilous species that may affect the availability of this breeding resource through interference or exploitation competition. In addition, we characterize body size, a key trait involved in competitive ability, for all *Nicrophorus* species at Whitehall Forest throughout the season. Finally, we compare our findings to other published natural history data for Nicrophorines. We document a significantly longer active season than was observed twenty years previously at Whitehall Forest for both *N. orbicollis* and *Nicrophorus tomentosus*, potentially due to climate change. As expected, the adult body size of *N. orbicollis* was larger than *N. tomentosus*, the only other *Nicrophorus* species that was captured in 2022 at Whitehall Forest. The other most prevalent interspecific insects captured included species in the families Staphylinidae, Histeridae, Scarabaeidae, and Elateridae, which may act as competitors or predators of *Nicrophorus* eggs and larvae. Together, our results indicate significant variation in intra- and interspecific competition relative to populations within the *N. orbicollis* range. These findings suggest that the competitive environment varies extensively over space and time, which help to inform the role of ecology in the evolution of parental care in this species.

## 1 INTRODUCTION

Burying beetles of the genus *Nicrophorus* (Silphidae) have long intrigued evolutionary biologists due to their complex parental care behavior (Pukowski, 1933; Eggert et al., 1997; Scott, 1998). Broadly, parental care is expected to evolve to ameliorate harsh environments for offspring (Wilson, 1975), and variation in parental care can be driven by ecological contexts that change the costs and benefits of allocating effort to current and future reproductive opportunities (Stearns, 1989; Richardson et al., 2020). The costs and benefits of parental care, and what strategies are pursued, are determined in part by competition for mates and breeding resources (Richardson et al., 2020). Moreover, parental care in *Nicrophorus* is known to influence development of offspring body size, a key trait essential for competition over breeding resources (Otronen, 1988; Hopwood et al., 2014; Lee et al., 2014; Jarrett et al., 2017). The breeding season of burying beetles can encompass multiple generations that experience different competitive environments depending on habitat and when individuals reach adulthood (DeMoss, 1968; Anderson, 1982; Scott et al., 1990; Eggert & Müller, 1997; Scott, 1998; Meierhofer et al., 1999). Thus, understanding the evolution of parental care in this system requires an understanding of how competition and ecological context vary over a breeding season in wild populations.

Here, we characterize the competitive environment and traits related to competition in *Nicrophorus orbicollis* across its breeding season in the southern portion of their range (**Figure 1**). *Nicrophorus orbicollis*, like all Nicrophorinae, require small vertebrate carcasses to breed, which are an ephemeral and highly sought-after resource. Body size is a key competitive trait for *Nicrophorus*, as larger beetles usually win contests for carcasses (Otronen, 1988; Robertson, 1993), and parental care itself has a strong influence on offspring development and adult body size (Hopwood et al., 2014; Jarrett et al., 2017). Parental care in *N. orbicollis* is thought to have evolved to buffer offspring from competition, predation by scavengers, and decomposition of the carcass by bacteria and fungi (Eggert & Müller, 1997; Scott, 1998). Once beetles have found a carcass, they strip the exterior (e.g., fur or feathers), bury, and maintain the carcass with an elaborate cocktail of secretions that minimize decomposition (Eggert & Müller, 1997; Scott, 1998; Arce et al., 2012). Following the hatch of larvae, parents then regurgitate pre-digested carrion to their begging larvae (Pukowski, 1933; Milne et al., 1976).

**Figure 1.**
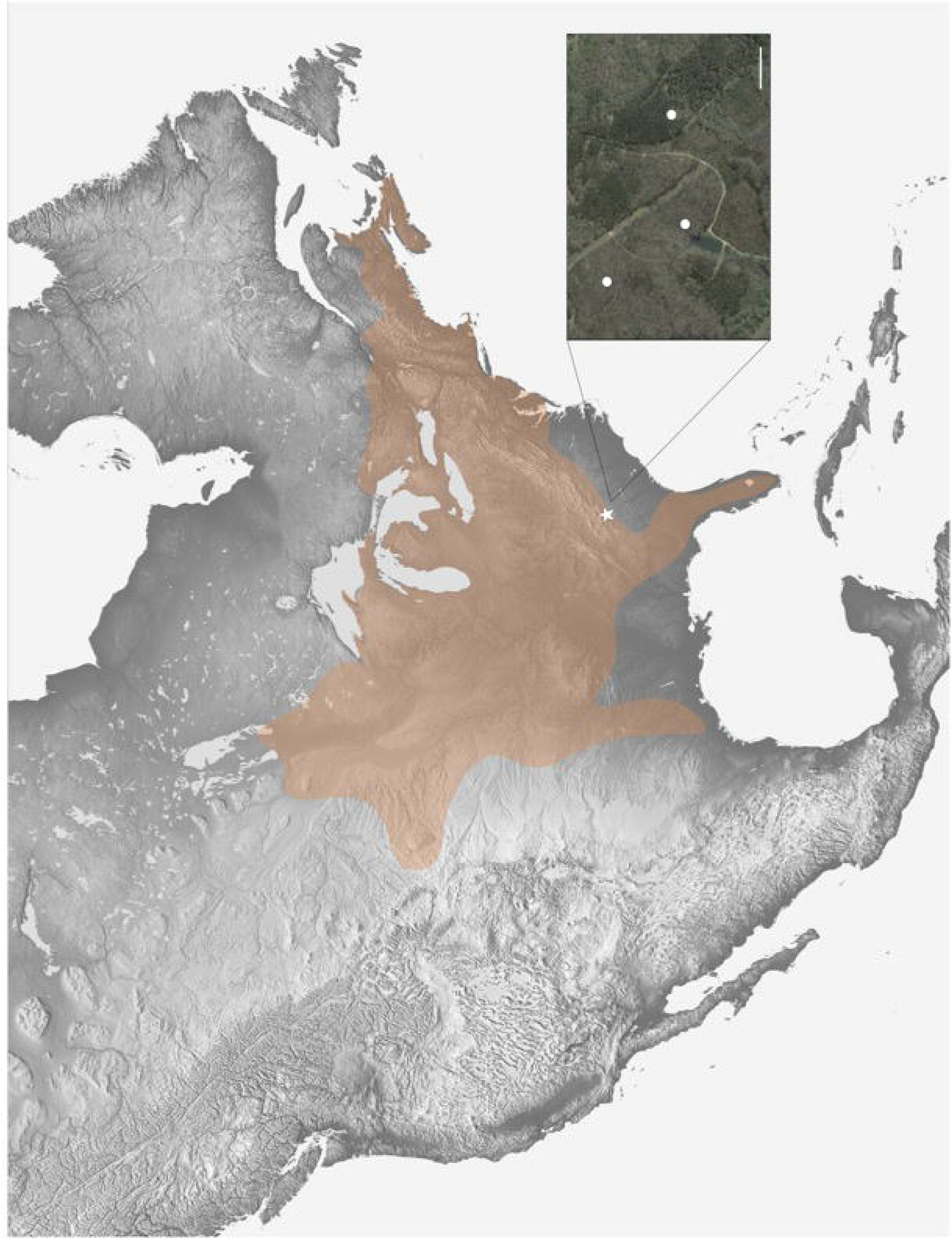
Range of *N. orbicollis* and location of study sites. Map derived from verified iNaturalist sightings indicating the general range of *N. orbicollis* (light orange overlay). Inset indicates relative location of study area within the *N. orbicollis* range and distribution of trap lines (white circles) within Whitehall Forest. Scale bar indicates 150 meters.

Burying beetle parents actively defend their carcass from intra- and interspecific intruders. These interactions can be influenced by temperature (Wilson et al., 1984), and parental care itself may be influenced by temperature (Meierhofer et al., 1999; Benowitz et al., 2019). Temperature affects both seasonal transitions between overwintering behavior and variation in the competitive environment. Insect species that use carrion show varying sensitivities to temperature and humidity, which likely influence the competitive environment (Wilson et al., 1984). *Nicrophorus* species show temporal niche partitioning during both circadian and seasonal cycles that are likely supported physiologically by differences in temperature tolerance (Anderson, 1982; Keller et al., 2019). Moreover, burial and prevention of decomposition may help to prevent discovery of the carcass by scavengers and competitors (Robertson, 1993; Pagh et al., 2015; Trumbo et al., 2021), particularly because decomposition is strongly affected by temperature. Outcomes of competition and population dynamics thus reflect a combination of species presence and their activity levels at different temperatures. For this reason, we collected temperature, phenological, and competition data to determine how the competitive environment changes over the breeding season.

## 2 MATERIALS AND METHODS

### 2.1 Range map and study area

During the spring – fall of 2022, we conducted a survey of *N. orbicollis* abundance and sympatric necrophilous insects at Whitehall Forest in Clarke County, Georgia, USA, located in the southern portion of the *N. orbicollis* range (**Figure 1;** 33.8848° N, 83.3577° W). There are no published range maps of *N. orbicollis* in the literature; thus, we constructed an approximate range map for *N. orbicollis* from a GBIF Occurrence Download using verified “research-grade” iNaturalist (inaturalist.org) observations occurring from 2000 – 2022, downloaded from GBIF.org site on October 17, 2022 (https://doi.org/10.15468/dl.dregg3). Based solely on these data we produced a map in Adobe® Photoshop (v. 20.0.8; https://adobe.com) to show the approximate distribution of *N. orbicollis*, although it is likely that *N. orbicollis* occupy habitat outside the scope of our range map. The *N. orbicollis* range was overlaid on a topographic map generated by NASA’s Shuttle Radar Topography Mission in 2000 (https://photojournal.jpl.nasa.gov/catalog/PIA03377) and was rendered to grayscale in Adobe Photoshop to enable clearer visualization of the range.

Whitehall Forest is a discontinuous stretch of forest managed by the University of Georgia Warnell School of Forestry and covers approximately 840 acres of an “island” of forest surrounded by residential development in the Southern Outer Piedmont (subregion of the Georgia Piedmont ecoregion). The forest is comprised primarily of natural pine, planted pine, upland hardwood, and bottomland hardwood. Within our specific trapping areas, the predominant tree species included red maple (*Acer rubrum*), American hornbeam (*Carpinus caroliniana*), flowering dogwood (*Cornus florida*), North American beech (*Fagus grandifolia*), yellow poplar (*Liriodendron tulipifera*), sweetgum (*Liquidambar styraciflua*), shortleaf pine (*Pinus echinata*), southern red oak (*Quercus falcata*), post oak (*Quercus stellata*), and white elm (*Ulmus americana*). Understory was minimal, and the forest was primarily characterized by large, well-dispersed trees with deep leaf litter. The study sites are adjacent to habitat including brushy pastures and grassy fields, and much of Whitehall Forest is interspersed with areas of maintained grassland or areas of intermittent prescribed burns (**Figure 1 inset**). Based on maps generated from the USDA Soil Survey, soil in the study area is primarily comprised of sandy clay loam, loamy sandy, and alluvial land (**SI Appendix 1**), similar to other sites supporting *N. orbicollis* (DeMoss, 1968). Humus content was also high due to a large amount of leaf litter across the entirety of the study site.

*Nicrophorus* species require small vertebrate carcasses such as small mammals, reptiles, and birds to breed. The study area at Whitehall Forest supports approximately ten small mammal species that may be suitable for *N. orbicollis* reproduction (*pers. comm*. Steven Castleberry). In addition to small mammals, there are approximately 105 migratory and resident passerine bird species in the study area that fall within the appropriate weight range used by *N. orbicollis*. Whitehall Forest is also within the range of multiple species of reptiles that may support *N. orbicollis* breeding, although the species composition is not well known.

### 2.2 Trapping and life history of *Nicrophorus orbicollis* in Georgia

Beetles were trapped using hanging Japanese beetle traps (hereafter, traps; item no. 227723, Trécé Inc., Adair, OK, USA) baited with one-inch square cubes of salmon along three transect lines of five traps each (**Figure 1 inset**). Traps were hung approximately one meter off the ground in a tree that shaded the trap. We started trapping on March 1 and defined the onset of the *N. orbicollis* active season as when the first *N. orbicollis* was captured. We determined the end of the active season as two consecutive trapping events with no *N. orbicollis* captures. One transect was adjacent to a small pond and perennial stream, while the other two transects did not have permanent water. *N. orbicollis* is primarily found in woodland areas (Anderson, 1982), therefore, all traps were placed within woodland habitat. Traps were checked every 6 – 9 days from March 1 to December 15, except for a few instances where traps could not be checked at this frequency and the bait was not replaced for 14 days (for a list of the trapping dates, see **Appendix 2**). In traps, the salmon bait was contained in a small plastic cup that greatly reduced desiccation and insect scavenger consumption of the bait. For this reason, even though bait was not replaced weekly on several occasions, it was still present and able to attract beetles, as adults of *Nicrophorus* species are known to preferentially eat late stage decomposing meat (e.g., Rodriguez et al., 1983; Dekeirsschieter et al., 2011; von Hoermann et al., 2013). We thus calculated the number of beetles captured as the number per trap day to account for variation in when traps were checked. We further noted any teneral beetles (recently eclosed adults that have not yet become fully melanized), as they provide an indication of recent breeding activity.

We visualized the length of the activity period of *N. orbicollis* over the field season with data from similar studies both temporally within Whitehall Forest, using data collected in the study area in 2002 (Ulyshen & Hanula, 2003), and spatially, by using data from the central (Kentucky; DeMoss, 1968) and northern (Ontario; Anderson, 1982) portions of the *N. orbicollis* range. While all three studies used different trapping methods, e.g., Anderson (1982) used pitfall traps baited with carrion, their data are roughly comparable to data collected for this study as they both show seasonal activity of *N. orbicollis*, as well as periods of peak activity. Data were readily available in a tabular form in DeMoss (1968) but raw data were collected from Figure 8 in Anderson (1982) and Figure 1 in Ulyshen and Hanula (2003) using the online platform for PlotDigitizer (https://plotdigitizer.com/). All three studies collected data weekly, although there were gaps in the collection times across studies. To allow for visualization across studies that differed in the date that traps were checked, we assigned a single date to trap dates that were most closely aligned (i.e., three or fewer days). To account for gaps in data collection at Whitehall Forest relative to other sites, we summed the number of beetles collected at other sites over the same time periods when traps in Georgia were not checked. See **Appendix 2** for details of total captures per study-specific trap date.

### 2.3 Characterization of the competitive environment

We characterized the competitive environment for *N. orbicollis* by identifying all species of insects attracted to carrion in traps. Furthermore, we identified all individuals captured to genus and to species, if possible (**Table 1**), apart from those that appeared two or fewer times, as these likely reflect incidental bycatch. A type specimen of each species was collected and pinned. We measured pronotum length of all *Nicrophorus* captured to the nearest 0.01 mm using digital calipers and identified the sex of each beetle. A small subset of the *N. orbicollis* captured were retained (n = 144) to augment our laboratory population. Beetles sometimes escaped after sexing but prior to measurement (N = 3) or were too damaged or desiccated, in which case, they were identified to species but no sex or pronotum length data were collected (N = 77). Since burying beetles can travel long distances to find carrion, e.g., *Nicrophorus americanus* can travel 7.24 km in a single night (Jurzenski et al., 2011), the retention of a small number of individuals is unlikely to significantly impact local populations or results of trapping efforts. We did not retain more than 10 individuals in a week as precaution. Also, it is likely that some individuals were recaptured. Despite this, our capture methods were intended to demonstrate the number of individuals competing for carrion at any given time rather than provide total numbers of individuals in the study site and are thus still appropriate and representative of changes in the competitive environment.

We collected ambient temperature and precipitation data from the University of Georgia Weather Network (College of Agricultural and Environmental Sciences, University of Georgia), using the Watkinsville-HORT Weather Station, located 5.15 km from the nearest trapping location. *N. orbicollis* are known to survive overwintering at soil depths between 5 cm – 105 cm in more northern portions of their range (Hoback et al., 2015). For this reason, we placed Thermochron^®^ iButton temperature loggers (©Maxim Integrated Products, Inc., San Jose, CA) approximately 10-12 cm underground at the start of each transect line in October to determine the soil temperature associated with *Nicrophorus* diapause in our study area.

### 2.4 Statistical analyses

We performed all statistical analyses using JMP Pro (v. 16.0.0, http://jmp.com) and produced figures in SigmaPlot (v. 14.5, http://www.sigmaplot.co.uk). Results are means ± SE unless otherwise noted. We analyzed pronotum length of individuals by sex and by species using pooled t-tests and two-way ANOVA with sex and species as factors.

## 3 RESULTS

### 3.1 Life history of *N. orbicollis* in Georgia

We observed the first *N. orbicollis* activity in traps on 24 March (1 adult male, 1 adult female; **Figure 2a**), and captured the last *N. orbicollis* on November 16, roughly corresponding to when soil temperatures 12 cm underground fell below 14°C (**Figure 2c**). We captured four teneral *N. orbicollis* adults on 28 June. Developmental periods for *N. orbicollis* in the laboratory are approximately 45 days from egg to adult (Potticary et al., in review). Thus, these data suggest that the parents of these individuals laid eggs between mid- to late-May.

**Figure 2.**
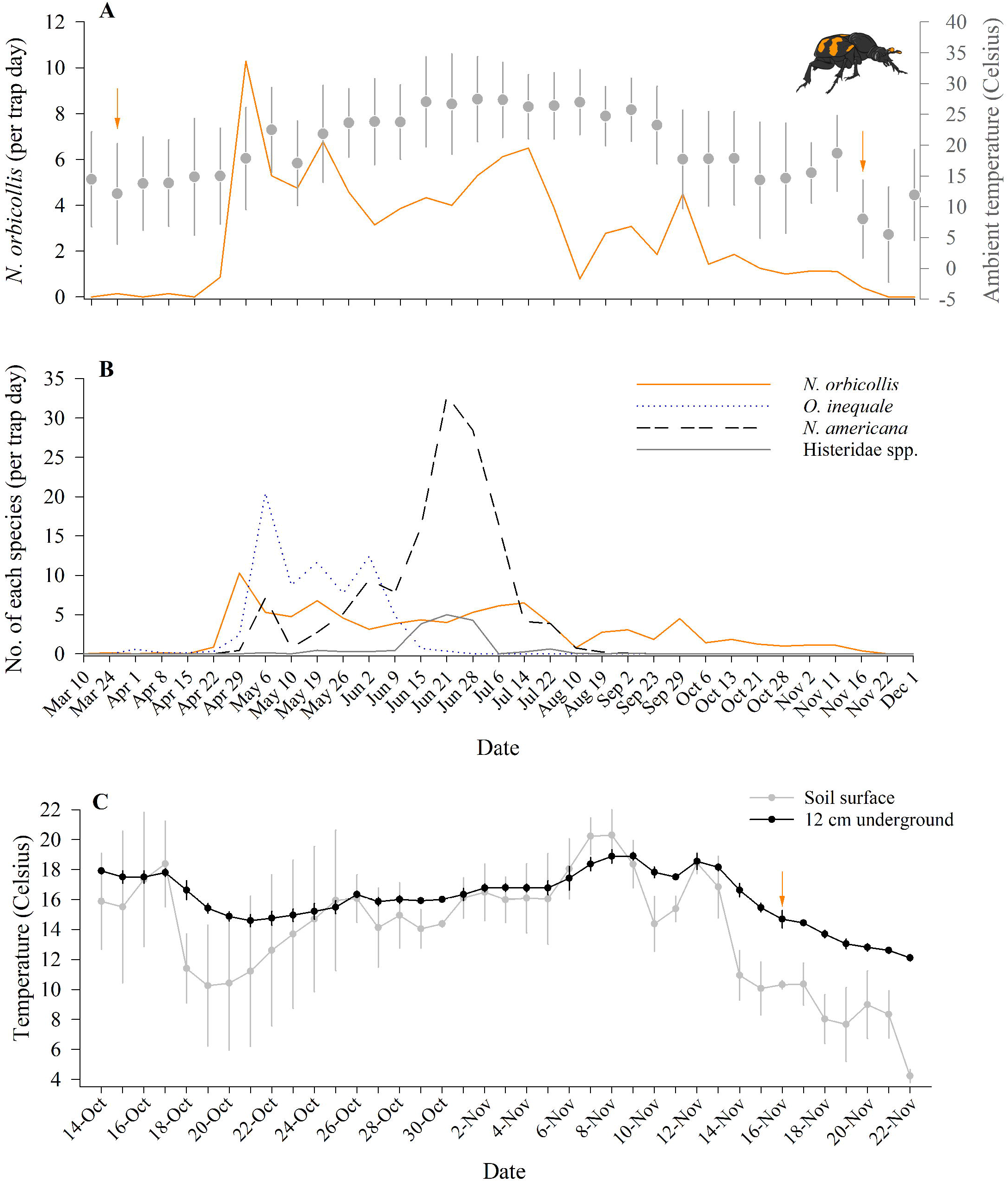
Activity of *N. orbicollis* relative to environmental context. (***A***) Number of *N. orbicollis* captured per trap day across the active season (orange line). Orange arrows indicate dates when the first or last *N. orbicollis* were captured for the season. Gray y-axis indicates mean ambient temperature (gray circle and error bars) from nearby weather station. (***B***) Number of each species per trap day for the three most abundant interspecific competitors (*Oiceoptoma inaequale*, *Necrophila americana*, and Histeridae spp.) for carrion relative to *N. orbicollis* captures. X-axis the same for both 2*A* and 2*B*. (C) Temperature at soil surface (gray line and gray circles) and 12 cm underground (black line and black circles) in study area. Note that temperature data from the study area is only available for October 14 – November 22, 2022. Error bars ± SD.

We observed an extension in the active season of *Nicrophorus* species in Whitehall Forest relative to research conducted in the same area in 2002 (**Figure 3**). *Nicrophorus orbicollis* emerged two weeks earlier and entered diapause nearly a week later, an approximately three-week extension of the active season over twenty years (**Figure 3a**). We observed a similar pattern in *N. tomentosus*, which emerged a week earlier and entered diapause two weeks later than in 2002 (**Figure 3b**). Across latitudes, *N. orbicollis* in more northern populations emerge later and enter diapause sooner than in Georgia, showing higher and more condensed bursts of activity (**Figure 4**). Together, these data support the idea that the length of the *N. orbicollis* active season tracks temperature both within and across populations.

**Figure 3.**
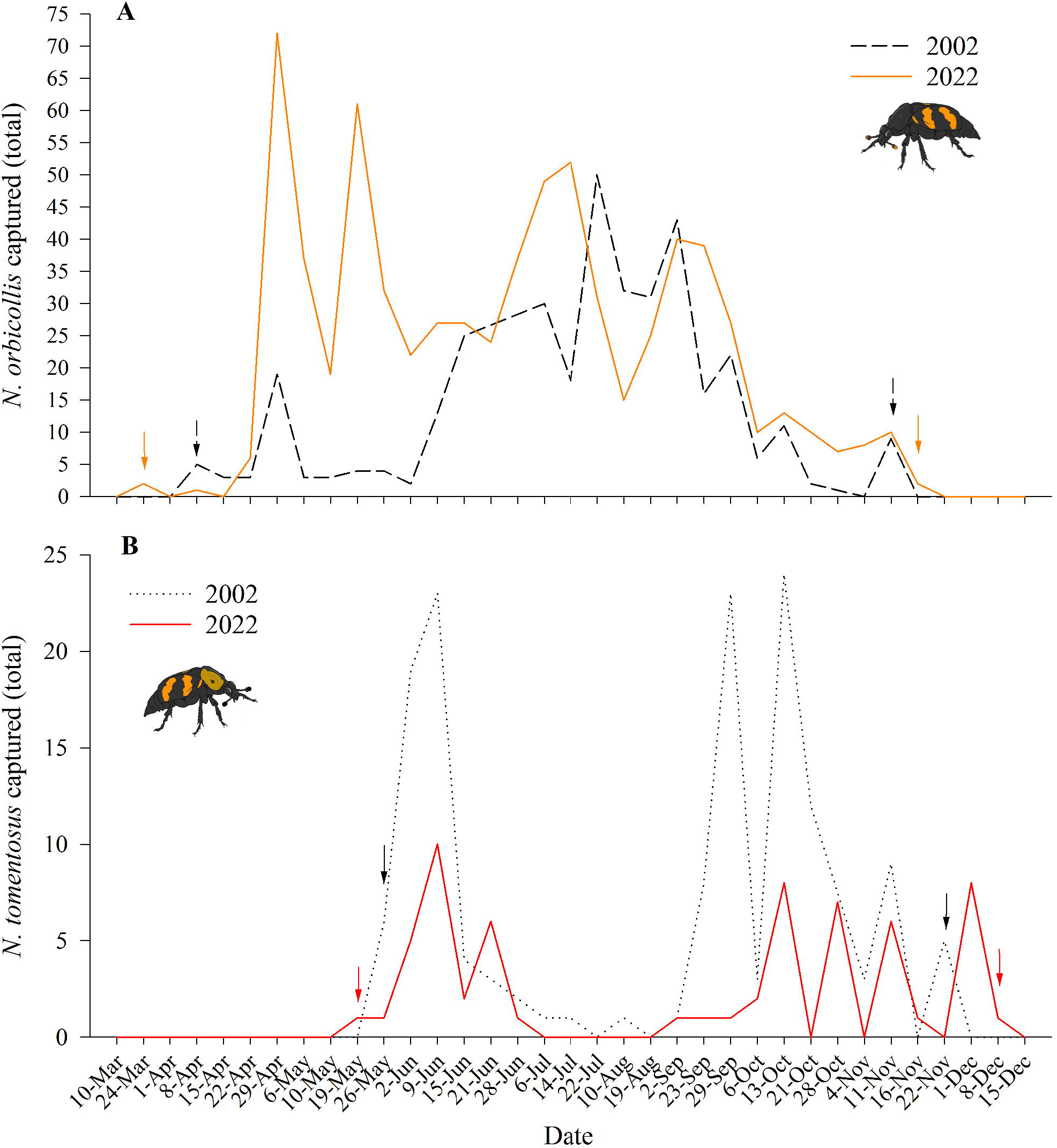
*Nicrophorus* active season has extended over twenty years. (***A***) *N. orbicollis* total captures at Whitehall Forest for 2002 (dashed black line) and 2022 (solid orange line). Black arrows indicate dates that *N. orbicollis* were first or last captured in 2002, while orange arrows indicate dates that *N. orbicollis* were first or last captured in 2022. (***B***) *N. tomentosus* total captures at Whitehall Forest for 2002 (dotted black line) and 2022 (solid red line). Black arrows indicate dates that *N. tomentosus* were first or last captured in 2002, while red arrows indicate dates that *N. tomentosus* were first or last captured in 2022. Note that scaling of y-axes for ***A*** and ***B*** differ. All 2002 data were acquired from (Ulyshen & Hanula, 2003).

**Figure 4.**
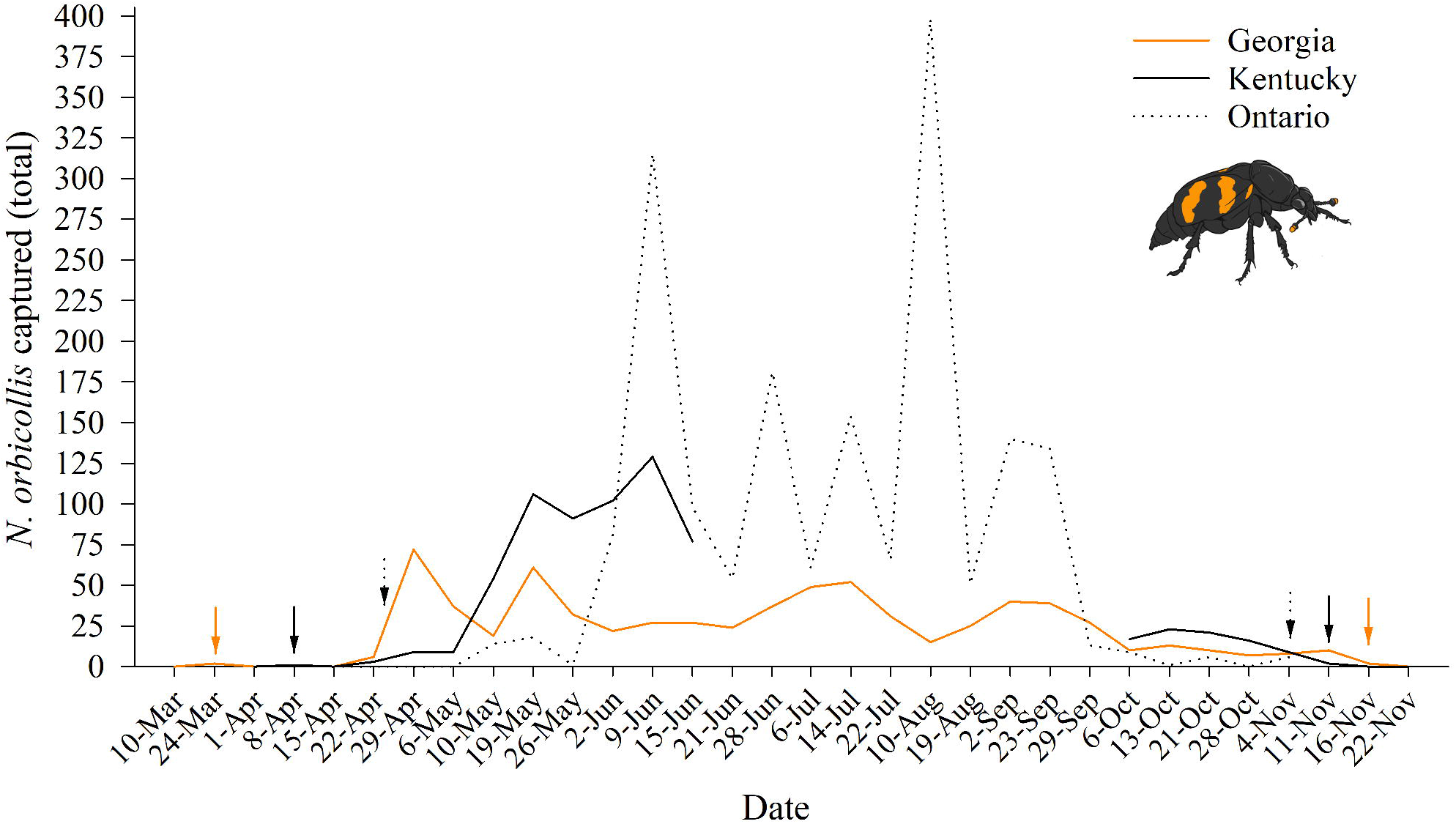
*N. orbicollis* active season longer in more southern areas. Weekly total *N. orbicollis* captures at Whitehall Forest, GA (solid orange line), in Kentucky (solid black line), and Ontario (dotted black line). Orange arrows indicate first and last *N. orbicollis* captures in Georgia, while solid black and dotted arrows indicate the same information for Kentucky (DeMoss, 1968) and Ontario (Anderson, 1982), respectively.

### 3.2 Characterization of the inter- and intraspecific competitive environment

A total of 703 *N. orbicollis* were captured over the season, with the number of females (N = 373) exceeding the numbers of males (N = 253; **Table 1**). A similar pattern was observed with *N. tomentosus*; a total of 53 of *N. tomentosus* were captured over the active season, with the number of females (N = 38) exceeding the number of males (N = 15). The most common interspecific species captured in carrion traps were *Necrophila americana*, *Oiceoptoma inaequale*, and Histeridae spp., although multiple potentially necrophilous species were captured (**Table 1**). The number of individuals of each species captured appears to coincide with a decrease in *N. orbicollis* captures and warmer ambient temperatures (**Figure 2b**). Peak periods of *N. tomentosus* captures appear to be disjunct from peak periods of *N. orbicollis* captures (**Figure 3**). The active season of *N. orbicollis* appears to be influenced on latitude, likely due to differences in temperature, with more northernly populations having a shorter active season than those farther south (**Figure 4**).

Pronotum length differed between *Nicrophorus* species and between the sexes in *N. tomentosus* (**Figure 5**). *N. orbicollis* was larger than *N. tomentosus* (*F*_1,673_ = 62.083, P < 0.0001), and there was an effect of sex on pronotum size (*F*_1,673_ = 5.240, P = 0.0224), but there was no interaction between sex and species on pronotum size (*F*_1,673_ = 2.873, P = 0.091). Pronotum length of female *N. orbicollis* ranged from 3.69 - 6.9 mm (mean 5.43 ± 0.03 mm, N = 371), while pronotum length of male *N. orbicollis* ranged from 3.65 – 6.73 mm (mean 5.37 ± 0.04 mm, N = 252). There was no statistical difference in the pronotum length of female versus male *N. orbicollis* (t_1,621_ = −1.1478, P = 0.252). Pronotum length of *N. tomentosus* females range from 3.4 – 5.96 mm (mean 4.83 ± 0.09 mm, N = 37), and male *N. tomentosus* body size ranged from 3.8 – 5.3 mm (mean 4.45 ± 0.15 mm, N = 15). On average, female *N. tomentosus* were larger than male *N. tomentosus* (t_1,50_ = −2.19, P = 0.030) but there was a large discrepancy in the sample size between males and females.

**Figure 5.**
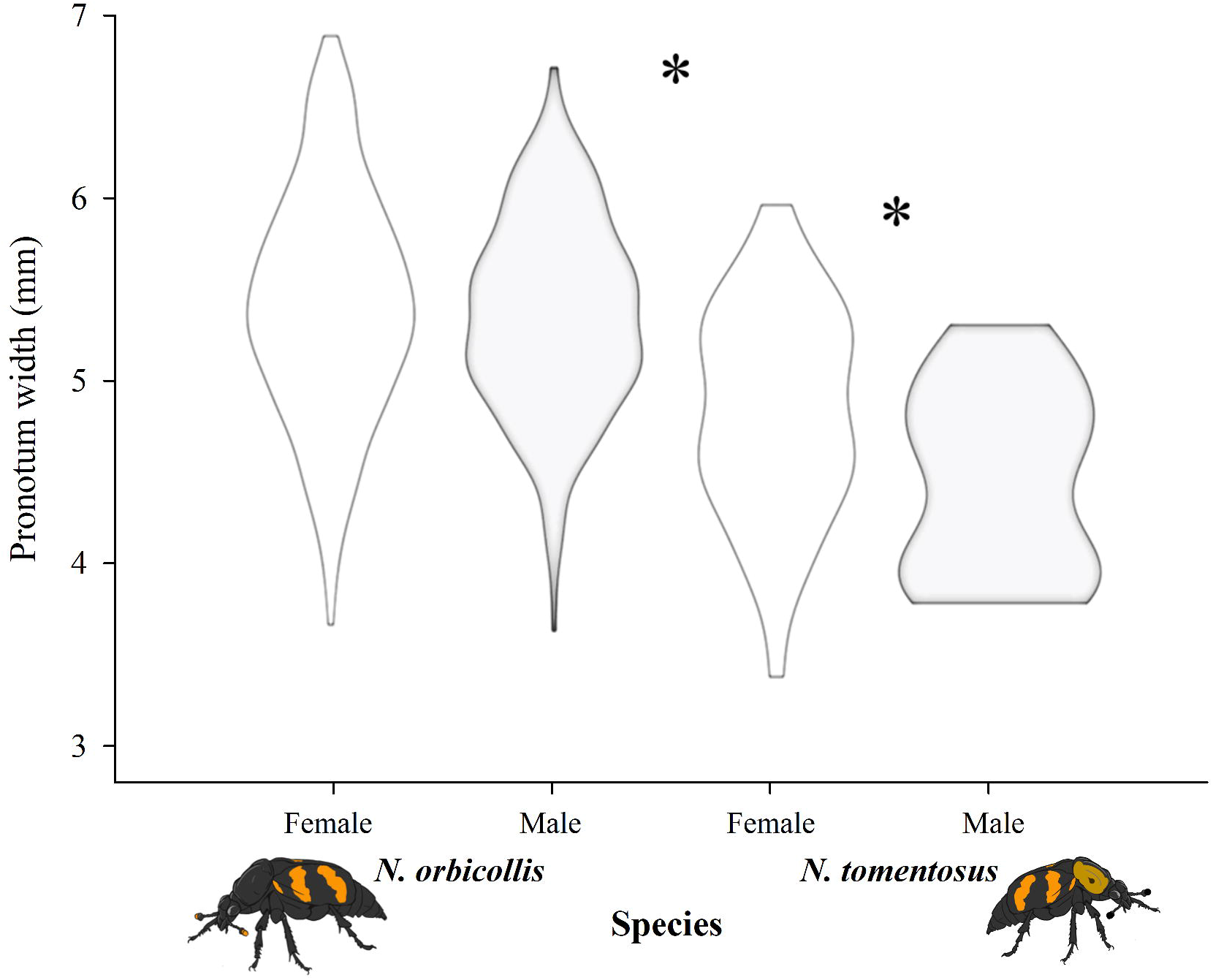
Variation in pronotum size of *Nicrophorus* at Whitehall Forest. Pronotum width of female and male *N. orbicollis* and *N. tomentosus*. While female (white violin) and male (grey violin) *N. orbicollis* do not differ in size, as measured by pronotum width, *N. orbicollis* is larger than *N. tomentosus*, and *N. tomentosus* females are larger than *N. tomentosus* males. Asterisks indicate a statistically significant difference between groups (P < 0.05).

## 4 DISCUSSION

We characterized the competitive environment across the active season for *N. orbicollis* breeding in a population located in southern portion of their range and compared these results to similar studies conducted at different temporal and spatial locations across their range. Of particular interest, we document that *Nicrophorus* species currently emerge earlier and enter diapause later in Whitehall Forest than 20 years ago (**Figure 3**). While this only constitutes two sampling periods, a difference of nearly three weeks (or about ½ the length of time to develop from egg to adult) in the length of the active season for two different species may indicate that climate change is impacting *Nicrophorus* activity, as warming temperatures have been indicated for Georgia (Frankson et al., 2022).

Insect diapause activity can be driven by photoperiod, density, and other factors (reviewed in Gill et al., 2017); however, *Nicrophorus* termination of diapause is likely driven by soil temperature as the beetles are thought to burrow to avoid winter temperatures and are thus unlikely to have access to other cues such as photoperiod (Hoback & Conley, 2015). Moreover, given the broad range for *N. orbicollis* (**Figure 1**), the most parsimonious explanation is that diapause activity is determined by local temperature cues (**Figure 4**). This may make *Nicrophorus* species particularly susceptible to changing temperatures resulting from climate change. The mechanisms that govern the duration of diapause and activity periods in *N. orbicollis* are poorly understood and warrant future research, and long-term data are particularly needed.

We found evidence indicating that *N. orbicollis* breeds as early as May at Whitehall Forest. Several teneral adults were observed on 28 June and given the approximately 45-day egg-to-adult developmental times in this species, this indicates an onset of breeding time between mid-May to late-May. Anderson (1982) used more in-depth methods to characterize tenerals in Ontario where they were not detected until August and September. A spike of young adults this close to the end of the breeding season (concluding in late September), may indicate that in more northern populations there is only a single generation per active season. An earlier onset of breeding in Whitehall Forest, in conjunction with delayed diapause relative to other portions of the range (Figure 4; DeMoss, 1968; Anderson, 1982), indicate that populations in Georgia may have multiple generations within a single season. Future research should investigate whether the expression of parental care varies in populations with single versus multiple generations per season.

The total number of *N. orbicollis* captured in this study was over double the number of individuals captured in Whitehall Forest in 2002, while there was nearly a 2/3 decrease in the number of *N. tomentosus* captured in 2022 relative to 2002 (Figure 3; Ulyshen & Hanula, 2003). While there are methodological differences between these studies – e.g., Ulyshen and Hanula (2003) killed and collected all individuals and baited traps with chicken – there are several lines of evidence suggesting that these differences reflect increases in *N. orbicollis* density. First, there is no evidence suggesting that either burying beetle species prefers chicken to salmon, or that the number of traps affected capture success, as similar numbers of *N. orbicollis* were captured for the last half of our field seasons (**Figure 3**). Second, our laboratory maintained the same trap lines in 2021 and collected 630 *N. orbicollis* to establish a laboratory colony, which also surpasses numbers captured in 2002. The significant decrease in the number of *N. tomentosus* documented between the two studies may also indirectly support that *N. orbicollis* populations are increasing. *N. tomentosus* are generally competitively subordinate to *N. orbicollis* (Schrempf et al., 2021), such that there is temporal niche partitioning between periods of activity of *N. orbicollis* and *N. tomentosus* (Anderson, 1982; Wilson et al., 1984; Keller et al., 2019). In accordance with previous research, we also found that *N. tomentosus* are more abundant in traps when *N. orbicollis* captures are low. An increase in *N. orbicollis* populations could provide one possible explanation for the drastic decrease in *N. tomentosus* at our study site. We suggest it would be valuable to use mark-recapture methods to derive estimates of population-level change over time in future research.

We captured more females than males of both *Nicrophorus* species. Across *Nicrophorus* species, some studies have documented roughly equal sex ratios in the wild (Milne & Milne, 1976; Anderson, 1982; Otronen, 1988), while others have also noted higher captures rates of females than males (Conley, 1982; Trumbo, 1990; Sikes, 1996). However, this is not necessarily indicative of differences in secondary sex ratios. Variation in captures may reflect differences in attraction to different types of carrion, such as state of decomposition, size, and species of carrion, as this may attract beetles at different reproductive stages (Wilson & Knollenberg, 1984; Otronen, 1988; Delclos et al., 2021). Males may not be as attracted to carcasses as females as they can “call” for females off a carcass and so males may split their time between searching for carcasses and calling for females (Müller et al., 1987; Eggert & Müller, 1997; Beeler et al., 1999). Moreover, our laboratory colony is derived from beetles at Whitehall Forest, and sex ratios of beetles that survive to adulthood are almost exactly 1:1 (Potticary et al., in review). This provides indirect evidence that the discrepancy in captures between the sexes may reflect behavioral differences in response to variation in carrion, although future research would be needed to address this effect.

*Nicrophorus orbicollis* densities were highest in late April – late May, and mid-June to early August (**Figure 3**). On a short timescale, the number of *Nicrophorus* that we captured reflect a combination of total individuals, temperature, food, and how many individuals are underground breeding and thus not searching for food. On an evolutionary timescale, we would expect peak *Nicrophorus* abundance to reflect an evolved response to ideal breeding conditions. In Georgia, April and May were periods of peak *N. orbicollis* capture, which correspond roughly to avian breeding in Whitehall Forest, and across latitudes, patterns are similar (**Figure 4**). *Nicrophorus orbicollis* depends on carcasses, which means that vertebrates dying of predation are unlikely to be available for breeding. Nestling and fledgling mortality of avian species due to exposure is high, and adults deposit dead nestlings away from the nest, which may provide a predictable source of carcasses for burying beetles. In addition, small mammal species are often plentiful in spring and early summer in North America (Merritt, 2010), and even though a reliable measure of mortality is difficult to obtain in some species, non-predator related mortality (e.g., food limitation, habitat conditions, disease) in larger populations could yield more carcass availability in spring. Together, these may provide a predicable source of breeding resources during certain months. Moreover, *N. orbicollis* captures in traps decreased in June, which would be consistent with beetles breeding on carcasses rather than searching for food (**Figure 2**). It is also intriguing that *N. tomentosus* is more cold tolerant and avoids *N. orbicollis*, yet *N. tomentosus* does not emerge from diapause until May despite low *N. orbicollis* numbers in March and early April. This may reflect that environmental conditions are too poor for successful breeding early in the season. However, future research is needed to determine the factors that influence peak activity periods across the *N. orbicollis* range.

While *Nicrophorus* are often expected to be the only species that compete via interference competition for breeding carcasses, we documented multiple species that may reduce the availability of carcasses through exploitation competition or that may be able to outcompete *Nicrophorus* due to larger body size. The most common inter-specific competitors captured, in order of abundance, included *Necrophila americana*, *Oiceoptoma inequale*, *N. tomentosus, Deltachilum gibbosum*, and *Oiceoptoma noveboracense* (**Table 1**). Some of these species are common across the range of *N. orbicollis*, albeit at different densities. For example, by-catch of research in Ontario found that the most common co-occurring species were *O. noveboracense*, *O. inequale*, and *N. americana* (Anderson, 1982). Intriguingly, previous work at Whitehall Forest recorded multiple species that were not captured or were only caught once in 2022, including *N. marginatus*, *N. pustulatus*, and *Necrodes surinamensis*. It is important to note that different trapping methods were used by Ulnysha and Hanula (2003) and *N. marginatus* and *N. pustulatus* were captured at low densities in 2002. Yet, these differences are unlikely to reflect a difference in trapping mode, as all three of these species were captured using similar trapping methods in Virginia (unpub. data). The absence of *N. surinamensis* is particularly stark, as this species was relatively common in 2002 and it was only captured a single time in 2022 in late November. It is possible that changes in species composition reflect stochasticity in sampling or, alternatively, that these differences reflect long-term changes in community composition due to other factors, like climate change. Climate change has been linked to changes in biological interactions that impact species composition across a diversity of systems (Dijkstra et al., 2011; Czortek et al., 2018). Our documentation of both an extended active season for two species of *Nicrophorus* species and density changes over the past twenty years at Whitehall may indicate effects of climate change.

We captured multiple species for which was unclear whether they were acting as competitors or predators of larvae on carcasses, including several species of the beetle families Elateridae and Histeridae (**Table 1**). Histerid beetles were one of the most abundant groups that we captured, although only one specimen was identified to genus (*Euspilota* sp.). Many histerid species are generalist predators that are known to prey upon small arthropods, including the immature and adult stages of other insects, and some histerid species are necrophilous (Correa et al., 2020). Histerid species are found worldwide and an estimated 6% are associated with carrion (Correa et al., 2020). Histeridae species have been captured in conjunction with *Nicrophorus* species in this study and other ecological studies all over the world (Shubeck, 1983; Naranjo-López et al., 2011; Psarev et al., 2020). Indeed, Naranjo-López and Navarrete-Heredia (2011) found that Histeridae species were one of the most abundant groups captured with *Nicrophorus olidus* in Mexico. Despite the widespread overlap of Histeridae and *Nicrophorus* species, there is little known about how hister beetles may impact breeding *Nicrophorus*. The threat of predation to offspring is expected to be a major factor in the evolution of parental care in insects like burying beetles (Tallamy, 1984; Scott, 1990, 1998; Suzuki et al., 2006; Trumbo, 2022), and thus widespread predators should be of importance for behavioral evolution across *Nicrophorus*. It is intriguing that the peak of hister beetle activity in this study corresponded to periods when breeding *N. orbicollis* likely had larvae on carcasses (early June to early July; **Figure 2**). Future research should investigate the relationship between Nicrophorus and associated histerid and elaterid species and examine any potential impacts on *Nicrophorus* parental care strategies.

In conclusion, we documented extreme variation in the competitive environment experienced by *N. orbicollis* over space and time. Behavior is a response to a context, and parental care strategies are expected to be particularly influenced by competition in burying beetle species. Thus, an individual burying beetle’s experience depends on when they are an adult and the population context of their natal population (Meierhofer et al., 1999). In this study, we document changes in both the length of the active season and the species composition of Whitehall Forest which may be influenced by climate change. Future behavioral and ecological research to better understand breeding strategies, activity, distribution, and effects of climate change on *N. orbicollis* is warranted.

## Acknowledgements

We would like to thank Kathryn Kollars for the beetle art used in this study. We would also like to thank Kathryn Kollars, Lance Fountain, and Emily Shelby for field assistance and help with preparation of insect collection. We would also like to thank Steven Castleberry and the mammalogy class at University of Georgia for providing the list of known small mammal species from within the study area. Lastly, we are indebted to the Warnell School of Ecology for allowing us to conduct this research at Whitehall Forest. ALP, AJM, and HWO conceived of study and developed methods. ALP and HWO collected data. ALP created figures. ALP, HWO, and JVM identified captured species. ALP, HWO, JVM, and AJM all contributed to editing and writing the manuscript. ALP was funded by a USDA cooperative agreement to AJM.

## Figures and Tables

**Table 1. Species captured in carrion traps across N. orbicollis active season.** Species for which at least ten were captured in carrion traps. Where possible, individuals were identified to species, although several were only identified to genus or family. For Nicrophorus species, individuals were also identified to sex, as described below the total capture numbers. Capture dates reflect the first and last dates individuals were captured for each.

