## Appendix 1 for "Nature Notes: Spatiotemporal variation in the competitive environment, with implications for how climate change may affect a species with parental care"

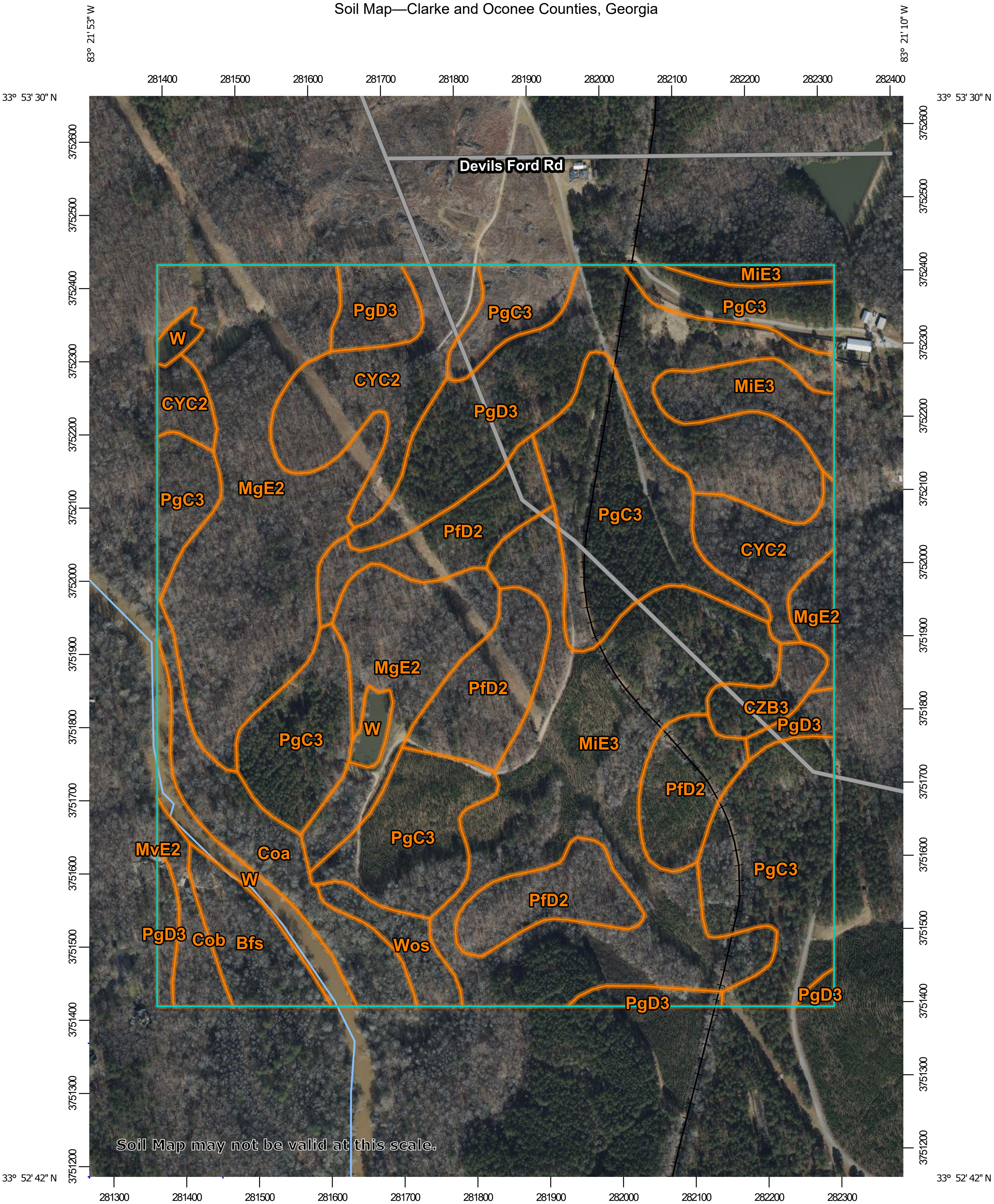


Map Scale: 1:7,210 if printed on A portrait (8.5" x 11") sheet.

Meters: 0, 100, 200, 400, 600

Feet: 0, 350, 700, 1400, 2100

Map projection: Web Mercator Corner coordinates: WGS84 Edge tics: UTM Zone 17N WGS84

#### MAP LEGEND

**Area of Interest (AOI)**
- Area of Interest (AOI)

**Soils**
- Soil Map Unit Polygons
- Soil Map Unit Lines
- Soil Map Unit Points

**Special Point Features**
- Blowout
- Borrow Pit
- Clay Spot
- Closed Depression
- Gravel Pit
- Gravelly Spot
- Landfill
- Lava Flow
- Marsh or swamp
- Mine or Quarry
- Miscellaneous Water
- Perennial Water
- Rock Outcrop
- Saline Spot
- Sandy Spot
- Severely Eroded Spot
- Sinkhole
- Slide or Slip
- Sodic Spot
- Spoil Area
- Stony Spot
- Very Stony Spot
- Wet Spot
- Other
- Special Line Features

**Water Features**
- Streams and Canals

**Transportation**
- Rails
- Interstate Highways
- US Routes
- Major Roads
- Local Roads

**Background**
- Aerial Photography

#### MAP INFORMATION

The soil surveys that comprise your AOI were mapped at 1:15,800.

Warning: Soil Map may not be valid at this scale.

Enlargement of maps beyond the scale of mapping can cause misunderstanding of the detail of mapping and accuracy of soil line placement. The maps do not show the small areas of contrasting soils that could have been shown at a more detailed scale.

Please rely on the bar scale on each map sheet for map measurements.

Source of Map: Natural Resources Conservation Service
Web Soil Survey URL:
Coordinate System: Web Mercator (EPSG:3857)

Maps from the Web Soil Survey are based on the Web Mercator projection, which preserves direction and shape but distorts distance and area. A projection that preserves area, such as the Albers equal-area conic projection, should be used if more accurate calculations of distance or area are required.

This product is generated from the USDA-NRCS certified data as of the version date(s) listed below.

Soil Survey Area: Clarke and Oconee Counties, Georgia
Survey Area Data: Version 15, Sep 14, 2022

Soil map units are labeled (as space allows) for map scales 1:50,000 or larger.

Date(s) aerial images were photographed: Dec 19, 2020—Dec 27, 2020

The orthophoto or other base map on which the soil lines were compiled and digitized probably differs from the background imagery displayed on these maps. As a result, some minor shifting of map unit boundaries may be evident.

### Map Unit Legend

| Map Unit Symbol | Map Unit Name | Acres in AOI | Percent of AOI |
|---|---|---|---|
| Bfs | Buncombe loamy sand | 5.1 | 2.2% |
| Coa | Congaree soils and alluvial land | 8.5 | 3.7% |
| Cob | Chewacla soils and alluvial land | 2.8 | 1.2% |
| CYC2 | Cecil sandy loam, 6 to 10 percent slopes, moderately eroded | 17.4 | 7.5% |
| CZB3 | Cecil sandy clay loam, 2 to 6 percent slopes, severely eroded | 2.7 | 1.2% |
| MgE2 | Madison sandy loam, 15 to 25 percent slopes, eroded | 40.5 | 17.4% |
| MiE3 | Madison sandy clay loam, 10 to 25 percent slopes, severely eroded | 41.7 | 17.9% |
| MvE2 | Musella clay loam, 15 to 25 percent slopes, eroded | 0.0 | 0.0% |
| PfD2 | Pacolet sandy loam, 10 to 15 percent slopes, moderately eroded | 23.0 | 9.8% |
| PgC3 | Pacolet sandy clay loam, 6 to 10 percent slopes, severely eroded | 53.0 | 22.7% |
| PgD3 | Pacolet sandy clay loam, 10 to 15 percent slopes, severely eroded | 30.7 | 13.1% |
| W | Water | 5.3 | 2.3% |
| Wos | Wehadkee and Alluvial land, wet | 2.8 | 1.2% |
| **Totals for Area of Interest** | | **233.5** | **100.0%** |
