## Appendix 2 for "Nature Notes: Spatiotemporal variation in the competitive environment, with implications for how climate change may affect a species with parental care"

| Site | Date beetles were collected from trap in paper | No. *N. orbicollis* captured | Revised date range for comparison across papers | Revised No. of *N. orbicollis* to account for missing dates |
|---|---|---|---|---|
| Whitehall 2022 | 10-Mar | 0 | 10-Mar | |
| Whitehall 2022 | 24-Mar | 2 | 24-Mar | |
| Kentucky 1968 | 29-Mar | 0 | 1-Apr | |
| Whitehall 2022 | 1-Apr | 0 | 1-Apr | |
| Kentucky 1968 | 6-Apr | 1 | 8-Apr | |
| Whitehall 2022 | 8-Apr | 1 | 8-Apr | |
| Ontario 1980 | 12-Apr | 0 | 15-Apr | |
| Kentucky 1968 | 12-Apr | 0 | 15-Apr | |
| Whitehall 2022 | 15-Apr | 0 | 15-Apr | |
| Kentucky 1968 | 19-Apr | 3 | 22-Apr | |
| Ontario | 19-Apr | 0 | 22-Apr | |
| Whitehall 2022 | 22-Apr | 6 | 22-Apr | |
| Ontario | 26-Apr | 0 | 29-Apr | |
| Kentucky 1968 | 26-Apr | 9 | 29-Apr | |
| Whitehall 2022 | 29-Apr | 72 | 29-Apr | |
| Kentucky 1968 | 3-May | 9 | 6-May | |
| Ontario | 6-May | 0 | 6-May | |
| Whitehall 2022 | 6-May | 37 | 6-May | |
| Whitehall 2022 | 10-May | 19 | 10-May | |
| Kentucky 1968 | 12-May | 54 | 10-May | |
| Ontario 1982 | 14-May | 14 | 10-May | |
| Kentucky 1968 | 17-May | 106 | 19-May | |
| Whitehall 2022 | 19-May | 61 | 19-May | |
| Ontario 1982 | 21-May | 18 | 19-May | |
| Kentucky 1968 | 25-May | 91 | 26-May | |
| Whitehall 2022 | 26-May | 32 | 26-May | |
| Ontario 1982 | 28-May | 1 | 26-May | |
| Kentucky 1968 | 31-May | 102 | 2-Jun | |
| Whitehall 2022 | 2-Jun | 22 | 2-Jun | |
| Ontario 1982 | 4-Jun | 81 | 2-Jun | |
| Kentucky 1968 | 8-Jun | 129 | 9-Jun | |
| Whitehall 2022 | 9-Jun | 27 | 9-Jun | |
| Ontario 1982 | 11-Jun | 315 | 9-Jun | |
| Kentucky 1968 | 15-Jun | 77 | 15-Jun | |
| Whitehall 2022 | 15-Jun | 27 | 15-Jun | |
| Ontario 1982 | 18-Jun | 98 | 15-Jun | |
| Whitehall 2022 | 21-Jun | 24 | 21-Jun | |
| Ontario 1982 | 25-Jun | 54 | 21-Jun | |
| Whitehall 2022 | 28-Jun | 37 | 28-Jun | |
| Ontario 1982 | 2-Jul | 181 | 28-Jun | |
| Whitehall 2022 | 6-Jul | 49 | 6-Jul | |

| | | | | |
|---|---|---|---|---|
| Ontario 1982 | 9-Jul | 61 | 6-Jul | |
| Whitehall 2022 | 14-Jul | 52 | 14-Jul | |
| Ontario 1982 | 16-Jul | 154 | 14-Jul | |
| Whitehall 2022 | 22-Jul | 31 | 22-Jul | |
| Ontario 1982 | 23-Jul | 66 | 22-Jul | |
| Ontario 1982 | 30-Jul | 170 | 10-Aug | Ontario 1982: 398 |
| Ontario 1982 | 6-Aug | 180 | 10-Aug | |
| Ontario 1982 | 13-Aug | 48 | 10-Aug | |
| Whitehall 2022 | 10-Aug | 15 | 10-Aug | |
| Whitehall 2022 | 19-Aug | 25 | 19-Aug | |
| Ontario 1982 | 20-Aug | 51 | 19-Aug | |
| Ontario 1982 | 27-Aug | 100 | 2-Sep | Ontario 1982: 140 |
| Whitehall 2022 | 2-Sep | 40 | 2-Sep | |
| Ontario 1982 | 3-Sep | 75 | 2-Sep | |
| Ontario 1982 | 10-Sep | 91 | 23-Sep | Ontario 1982: 134 |
| Ontario 1982 | 15-Sep | 24 | 23-Sep | |
| Ontario 1982 | 22-Sep | 19 | 23-Sep | |
| Whitehall 2022 | 23-Sep | 39 | 23-Sep | |
| Ontario 1982 | 29-Sep | 13 | 29-Sep | |
| Whitehall 2022 | 29-Sep | 27 | 29-Sep | |
| Kentucky 1968 | 4-Oct | 17 | 6-Oct | |
| Ontario 1982 | 5-Oct | 9 | 6-Oct | |
| Whitehall 2022 | 6-Oct | 10 | 6-Oct | |
| Kentucky 1968 | 11-Oct | 23 | 13-Oct | |
| Ontario 1982 | 13-Oct | 1 | 13-Oct | |
| Whitehall 2022 | 13-Oct | 13 | 13-Oct | |
| Kentucky 1968 | 18-Oct | 21 | 21-Oct | |
| Ontario 1982 | 20-Oct | 6 | 21-Oct | |
| Whitehall 2022 | 21-Oct | 10 | 21-Oct | |
| Kentucky 1968 | 25-Oct | 16 | 28-Oct | |
| Ontario 1982 | 27-Oct | 0 | 28-Oct | |
| Whitehall 2022 | 28-Oct | 7 | 28-Oct | |
| Kentucky 1968 | 1-Nov | 9 | 4-Nov | |
| Whitehall 2022 | 2-Nov | 8 | 4-Nov | |
| Ontario 1982 | 3-Nov | 6 | 4-Nov | |
| Kentucky 1968 | 8-Nov | 2 | 11-Nov | |
| Whitehall 2022 | 11-Nov | 10 | 11-Nov | |
| Kentucky 1968 | 15-Nov | 0 | 16-Nov | |
| Whitehall 2022 | 16-Nov | 2 | 16-Nov | |
| Kentucky 1968 | 22-Nov | 0 | 22-Nov | |
| Whitehall 2022 | 22-Nov | 0 | 22-Nov | |

| Site | year | Date beetles were collected from trap in paper | No. *N. orbicollis* captured | No. *N. tomentosus* captured | Revised date range for comparison across papers | Revised No. of *N. orbicollis* to account for missing dates | Revised No. of *N. tomentosus* to account for missing dates |
|---|---|---|---|---|---|---|---|
| Whitehall | 2002 | 3-Jan | 0 | 0 | 3-Jan | 0 | 0 |
| Whitehall | 2002 | 3-Feb | 0 | 0 | 3-Feb | 0 | 0 |
| Whitehall | 2002 | 10-Feb | 0 | 0 | 10-Feb | 0 | 0 |
| Whitehall | 2002 | 17-Feb | 0 | 0 | 17-Feb | 0 | 0 |
| Whitehall | 2002 | 24-Feb | 0 | 0 | 24-Feb | 0 | 0 |
| Whitehall | 2002 | 3-Mar | 0 | 0 | 3-Mar | 0 | 0 |
| Whitehall | 2002 | 10-Mar | 0 | 0 | 10-Mar | 0 | 0 |
| Whitehall | 2022 | 10-Mar | 0 | 0 | 10-Mar | 0 | 0 |
| Whitehall | 2022 | 24-Mar | 2 | 0 | 24-Mar | 2 | 0 |
| Whitehall | 2002 | 31-Mar | 0 | 0 | 1-Apr | 0 | 0 |
| Whitehall | 2022 | 1-Apr | 0 | 0 | 1-Apr | 0 | 0 |
| Whitehall | 2002 | 7-Apr | 5 | 0 | 8-Apr | 5 | 0 |
| Whitehall | 2022 | 8-Apr | 1 | 0 | 8-Apr | 1 | 0 |
| Whitehall | 2002 | 14-Apr | 3 | 0 | 15-Apr | 3 | 0 |
| Whitehall | 2022 | 15-Apr | 0 | 0 | 15-Apr | 0 | 0 |
| Whitehall | 2002 | 21-Apr | 3 | 0 | 22-Apr | 3 | 0 |
| Whitehall | 2022 | 22-Apr | 6 | 0 | 22-Apr | 6 | 0 |
| Whitehall | 2002 | 28-Apr | 19 | 0 | 29-Apr | 19 | 0 |
| Whitehall | 2022 | 29-Apr | 72 | 0 | 29-Apr | 72 | 0 |
| Whitehall | 2002 | 5-May | 3 | 0 | 6-May | 3 | 0 |
| Whitehall | 2022 | 6-May | 37 | 1 | 6-May | 37 | 1 |
| Whitehall | 2002 | 12-May | 3 | 0 | 10-May | 3 | 0 |
| Whitehall | 2022 | 10-May | 19 | 0 | 10-May | 19 | 0 |
| Whitehall | 2002 | 19-May | 4 | 0 | 19-May | 4 | 0 |
| Whitehall | 2022 | 19-May | 61 | 1 | 19-May | 61 | 1 |
| Whitehall | 2002 | 26-May | 4 | 6 | 26-May | 4 | 6 |
| Whitehall | 2022 | 26-May | 32 | 1 | 26-May | 32 | 1 |
| Whitehall | 2002 | 3-Jun | 2 | 19 | 2-Jun | 2 | 19 |
| Whitehall | 2022 | 2-Jun | 22 | 5 | 2-Jun | 22 | 5 |

| | | | | | | | |
|---|---|---|---|---|---|---|---|
| Whitehall | 2002 | 10-Jun | 13 | 23 | 9-Jun | 13 | 23 |
| Whitehall | 2022 | 9-Jun | 27 | 10 | 9-Jun | 27 | 10 |
| Whitehall | 2002 | 17-Jun | 25 | 4 | 15-Jun | 25 | 4 |
| Whitehall | 2022 | 15-Jun | 27 | 2 | 15-Jun | 27 | 2 |
| Whitehall | 2022 | 21-Jun | 24 | 6 | 21-Jun | 24 | 6 |
| Whitehall | 2022 | 28-Jun | 37 | 1 | 28-Jun | 37 | 1 |
| Whitehall | 2002 | 8-Jul | 30 | 1 | 6-Jul | 30 | 1 |
| Whitehall | 2022 | 6-Jul | 49 | 0 | 6-Jul | 49 | 0 |
| Whitehall | 2002 | 15-Jul | 18 | 1 | 14-Jul | 18 | 1 |
| Whitehall | 2022 | 14-Jul | 52 | 0 | 14-Jul | 52 | 0 |
| Whitehall | 2002 | 22-Jul | 50 | 0 | 22-Jul | 50 | 0 |
| Whitehall | 2022 | 22-Jul | 31 | 0 | 22-Jul | 31 | 0 |
| Whitehall | 2002 | 29-Jul | 17 | 1 | 10-Aug | 32 | 1 |
| Whitehall | 2002 | 5-Aug | 15 | 0 | 10-Aug | | |
| Whitehall | 2022 | 10-Aug | 15 | 0 | 10-Aug | 15 | 0 |
| Whitehall | 2002 | 12-Aug | 10 | 0 | 19-Aug | 31 | 0 |
| Whitehall | 2002 | 19-Aug | 21 | 0 | 19-Aug | | |
| Whitehall | 2022 | 19-Aug | 25 | 0 | 19-Aug | 25 | 0 |
| Whitehall | 2002 | 26-Aug | 29 | 0 | 2-Sep | 43 | 1 |
| Whitehall | 2002 | 2-Sep | 14 | 1 | 2-Sep | | |
| Whitehall | 2022 | 2-Sep | 40 | 1 | 2-Sep | 40 | 1 |
| Whitehall | 2002 | 16-Sep | 10 | 4 | 23-Sep | 16 | 8 |
| Whitehall | 2002 | 23-Sep | 6 | 4 | 23-Sep | | |
| Whitehall | 2022 | 23-Sep | 39 | 1 | 23-Sep | 39 | 1 |
| Whitehall | 2002 | 30-Sep | 22 | 23 | 29-Sep | 22 | 23 |
| Whitehall | 2022 | 29-Sep | 27 | 1 | 29-Sep | 27 | 1 |
| Whitehall | 2002 | 6-Oct | 6 | 3 | 6-Oct | 6 | 3 |
| Whitehall | 2022 | 6-Oct | 10 | 2 | 6-Oct | 10 | 2 |
| Whitehall | 2002 | 14-Oct | 11 | 24 | 13-Oct | 11 | 24 |
| Whitehall | 2022 | 13-Oct | 13 | 8 | 13-Oct | 13 | 8 |
| Whitehall | 2002 | 21-Oct | 2 | 12 | 21-Oct | 2 | 12 |
| Whitehall | 2022 | 21-Oct | 10 | 0 | 21-Oct | 10 | 0 |
| Whitehall | 2022 | 28-Oct | 7 | 7 | 28-Oct | 7 | 7 |

| | | | | | | | |
|---|---|---|---|---|---|---|---|
| Whitehall | 2002 | 4-Nov | 0 | 3 | 4-Nov | 0 | 3 |
| Whitehall | 2022 | 2-Nov | 8 | 0 | 4-Nov | 8 | 0 |
| Whitehall | 2002 | 11-Nov | 9 | 9 | 11-Nov | 9 | 9 |
| Whitehall | 2022 | 11-Nov | 10 | 6 | 11-Nov | 10 | 6 |
| Whitehall | 2002 | 18-Nov | 0 | 0 | 16-Nov | 0 | 0 |
| Whitehall | 2022 | 16-Nov | 2 | 1 | 16-Nov | 2 | 1 |
| Whitehall | 2002 | 25-Nov | 0 | 5 | 22-Nov | 0 | 5 |
| Whitehall | 2022 | 22-Nov | 0 | 0 | 22-Nov | 0 | 0 |
| Whitehall | 2002 | 1-Dec | 0 | 0 | 1-Dec | 0 | 0 |
| Whitehall | 2022 | 1-Dec | 0 | 8 | 1-Dec | 0 | 8 |
| Whitehall | 2002 | 9-Dec | 0 | 0 | 8-Dec | 0 | 0 |
| Whitehall | 2022 | 8-Dec | 0 | 1 | 8-Dec | 0 | 1 |
| Whitehall | 2002 | 16-Dec | 0 | 0 | 15-Dec | 0 | 0 |
| Whitehall | 2022 | 15-Dec | 0 | 0 | 15-Dec | 0 | 0 |
